# Structural Determinants of Cholesterol Recognition in Helical Integral Membrane Proteins

**DOI:** 10.1101/2020.06.15.152025

**Authors:** B. Marlow, G. Kuenze, B. Li, C. Sanders, J. Meiler

## Abstract

Cholesterol (CLR) is an integral component of mammalian membranes. It has been shown to modulate membrane dynamics and alter integral membrane protein (IMP) function. However, understanding the molecular mechanisms of these processes is complicated by limited and conflicting structural data: Specifically, in co-crystal structures of CLR-IMP complexes it is difficult to distinguish specific and biologically relevant CLR-IMP interactions from a nonspecific association captured by the crystallization process. The only widely recognized search algorithm for CLR-IMP interaction sites is sequence-based, i.e. searching for the so-called ‘CRAC’ or ‘CARC’ motifs. While these motifs are present in numerous IMPs, there is inconclusive evidence to support their necessity or sufficiency for CLR binding. Here we leverage the increasing number of experimental CLR-IMP structures to systematically analyze putative interaction sites based on their spatial arrangement and evolutionary conservation. From this analysis we create three-dimensional representations of general CLR interaction sites that form clusters across multiple IMP classes and classify them as being either specific or nonspecific. Information gleaned from our characterization will eventually enable a structure-based approach for prediction and design of CLR-IMP interaction sites.

**SIGNIFICANCE:** CLR plays an important role in composition and function of membranes and often surrounds and interacts with IMPs. It is a daunting challenge to disentangle CLRs dual roles as a direct modulator of IMP function through binding or indirect actor as a modulator of membrane plasticity. Only recently studies have delved into characterizing specific CLR-IMP interactions. We build on this previous work by using a combination of structural and evolutionary characteristics to distinguish specific from nonspecific CLR interaction sites. Understanding how CLR interacts with IMPs will underpin future development towards detecting and engineering CLR-IMP interaction sites.

## INTRODUCTION

Lipid composition is important for integral membrane protein (IMP) incorporation, localization, and trafficking. Many substrates of IMPs are membrane-associated. Thus, it is imperative that we analyze IMP function with consideration of the lipid composition of the respective membrane. There is a large degree of diversity in lipids(1,2), which ranges from the chemical nature of the head group to the saturation and length of the acyl chains. For example, sphingosine-1-phosphate (S1P) is an intracellular signaling molecule that consist of a sphingosine backbone with a position 1 attached phosphate group. Degradation of S1P to its metabolite hexadecenal is linked to the pathogenesis of Sjögren-Larsson syndrome(3).

Sterols are a distinct class of lipids, which are synthesized via the mevalonate pathway(4). Sterols are defined by their tetracyclic structure (see Figure 1) but have differing modifications to the tail and head groups. For example, stigmasterol in plants contains C24 alkyl groups or ergosterol in fungi has additional double bonds in ring B(5). For this study we focus on the mammalian sterol cholesterol (CLR), not just because of its critical importance in humans but also because it has the most atomic-detail structural information available in the Protein Data Bank (PDB)(6). CLR is found at very high concentrations, up to ~40 mol%(7,8), in some mammalian cell membranes. It is a polycyclic amphipathic molecule that contains a small polar head group (−OH) mounted on an asymmetric sterane backbone(9,10). One face of the molecule is characterized as planar and denoted as the α-face, while the β-face has two methyl groups pointing outward from the plane in an orthogonal fashion(11) (see Figure 1). CLR in the plasma membrane generally appears to have no preference for one leaflet over the other, although some evidence to the contrary exists(12–14). While the mitochondrial membrane has a lower CLR content (10%), there is a clear bias for CLR in the outer mitochondrial membrane. Trafficking CLR to the inner mitochondrial membrane is in fact the rate-limiting step in the steroidogenesis biosynthesis pathway(15). CLR has often been studied in conjunction with membrane dynamics. Its presence alters the physical properties of the membrane by stiffening disordered phases in which it is present, sometimes even provoking a transition into the liquid-ordered phase(9,16) and thereby reducing the liquid-disordered phase(16,17). It has been hypothesized that CLR contributes to the formation of raft-like regions within membranes(18,19), although the concept and/or precise definition of ‘lipid-rafts’ remains controversial(20–22).

**Figure 1.**
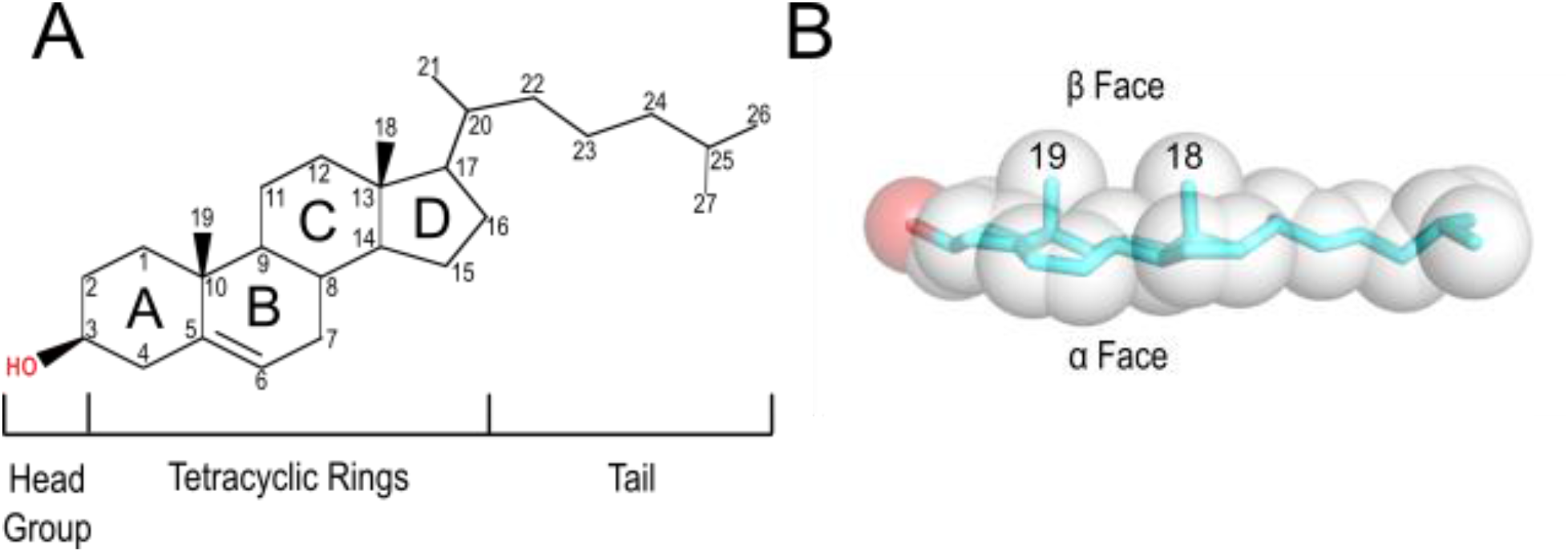
Structure of cholesterol. A. Chemical structure of cholesterol divided into the segments we used to calculate asymmetric preference. B. Space-filling and stick representation of cholesterol.

In recent years there have been several studies targeted towards understanding how CLR directly binds to IMPs and regulates their function. For example, CLR seems to preferentially bind to the inactive state of rhodopsin(23,24) or to inhibit the conductance of the Ca^2+^-sensitive K^+^ channels (BK)(25,26). CLR binding also suppresses the endothelial Kir2.1 channel that impairs flow-induced vasodilation and augments the development of atherosclerosis(27,28).

Innovative experiments have used CLR stereoisomers to distinguish between specific and nonspecific interactions(29–31). These analogues replicate several physiochemical membrane properties of CLR that would be consistent with modulating the membrane itself and thus nonspecific interactions with IMPs but would be incompatible with specific CLR-IMP interactions. If functional modification of an IMP by CLR is due solely from the impact of bulk CLR membrane properties, then there should be negligible differences in functional effects between natural CLR and its stereoisomers(32,33). For example, Levitan and Barbera(34) investigated the role of CLR’s stereospecificity and spatial orientation for binding to the Kir2.2 channel. A combination of molecular dynamics (MD) simulations, docking, electrophysiology and mutational studies uncovered that CLR, *epi*-CLR, and *ent*-CLR can all bind to the same pocket, but with different functions and predicted orientations(34,35). This study and others found that current predictive methods to identify CLR binding sites in IMPs yield mixed results at best(35). Characterization of possible CLR binding pockets at atomic detail is underdeveloped and hampers our understanding of CLR function.

For many years, efforts have been devoted to finding a consensus binding site on the surface of IMPs to differentiate between CLR as a solvent lipid and as a specific IMP ligand. Most prominently, the CLR recognition amino acid consensus (CRAC) sequence motif was proposed based on the Translocator Protein (TSPO) over 30 years ago(36). The CRAC motif is defined (from the N-to C-terminus with one-letter amino acid codes): (L/V)-X^1-5^-(Y)-X^1-5^-(K/R), where X^1-5^ represents one to five of any of the 20 amino acids. Later, an additional consensus sequence was proposed that is effectively an inverted-CRAC motif known as CARC, which is defined as (K/R)-X^1-5^-(Y/F)-X^1-5^-(L/V)(9,37,38). While, both motifs have been identified on multiple IMPs, there is inconclusive evidence of its correlation with known functional CLR binding sites(39). We could not find an explanation why phenylalanine is only required for the CARC motif and tryptophan is not required for either.

In 2013 Song et al identified 91 and 97 occurrences of the CRAC and CARC motifs in a dataset of 19 then-available co-crystal structures of CLR-IMP complexes respectively(39). Only one of the motifs was in the vicinity of a CLR binding site. It therefore remains unclear if the CRAC or the CARC motifs is sufficient and/or necessary for predicting specific CLR-IMP interaction sites. Since the Song paper was published, the number of CLR-IMP complexes has increased to 120 structures. For this study, we curated a dataset of 57 CLR-IMP complexes selecting high-resolution structures for which there is supporting evidence in the literature that the CLR-IMP interaction has biological relevance.

Here, we characterize the structural determinants of CLR recognition in 44 non-redundant CLR-IMP interaction sites and examine the results considering the locations of CRAC and CARC motifs. We clustered recurring modes of CLR binding into three structurally distinct classes and rank them as specific or nonspecific interaction sites based on their rate of evolution. We expect, that while this set is likely to be incomplete, it will provide a useful knowledgebase that can be expanded while simultaneously being used for prediction and design of CLR-IMP interaction sites. From this study we were able to categorize an interaction sites as specific or nonspecific based on the face of CLR interacting with the residues of the IMP.

## RESULTS

Our multistage analysis included both a low-resolution (amino acid composition) and high-resolution (atomic detail tertiary structure) investigation of the structural determinants for CLR-IMP interaction sites: 1. Assemble a dataset of high-resolution CLR-IMP complex co-crystal structures from the PDB. 2. Quantify the CRAC and CARC motifs involved in CLR recognition. 3. Breakdown the spatial arrangement of residues contacting the six segments of the CLR molecule (see Figure 1). 4. Find similarities between IMP families by clustering interaction sites. 5. Differentiate between specific and nonspecific CLR interaction sites based on a reduced rate of evolutionary drift. 6. Compare CLR-IMP interactions to specific CLR-soluble protein (SP) interaction sites.

### A CLR-IMP complex dataset was assembled

We created a dataset of experimentally determined high-resolution CLR-IMP structures that includes 57 IMPs from the PDB (see Table 1) as described in the Methods section. The resolution of the structures included in the analysis ranged from 1.5 Å to 3.5 Å. A distance cutoff of 6 Å between CLR and any non-hydrogen atoms on the IMP was used to define residues as part of the interaction site. Several structures included multiple CLR molecules resulting in 94 distinct CLR-IMP interaction sites. Next, IMP homologues or point mutants were excluded from the analysis to avoid double-counting, resulting in a set of 44 non-redundant IMP interaction sites (**bold** in Table 1). We hypothesize that there is a distinct difference between specific and nonspecific contacts made in these CLR-IMP interaction sites.

**Table 1.**
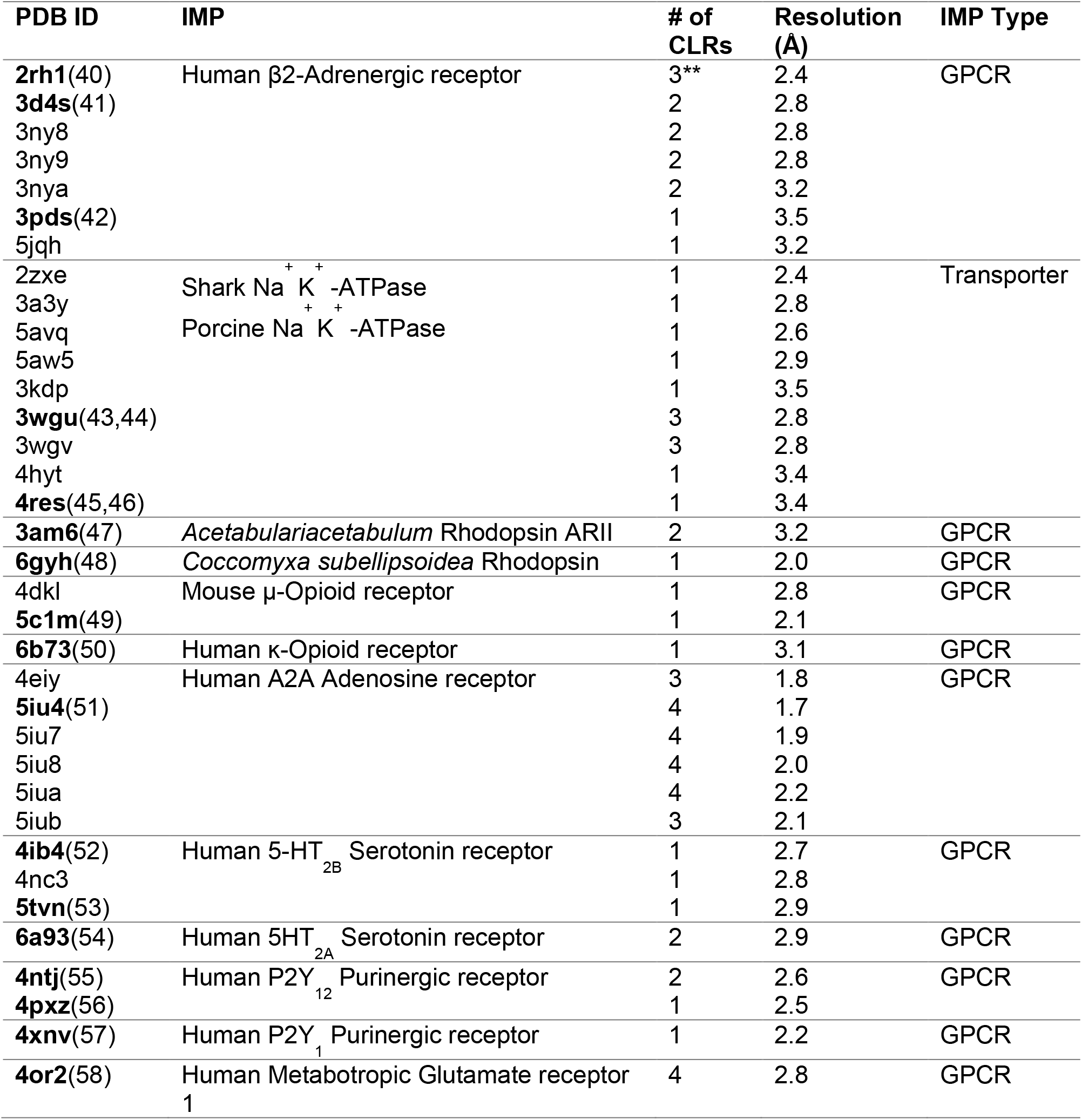
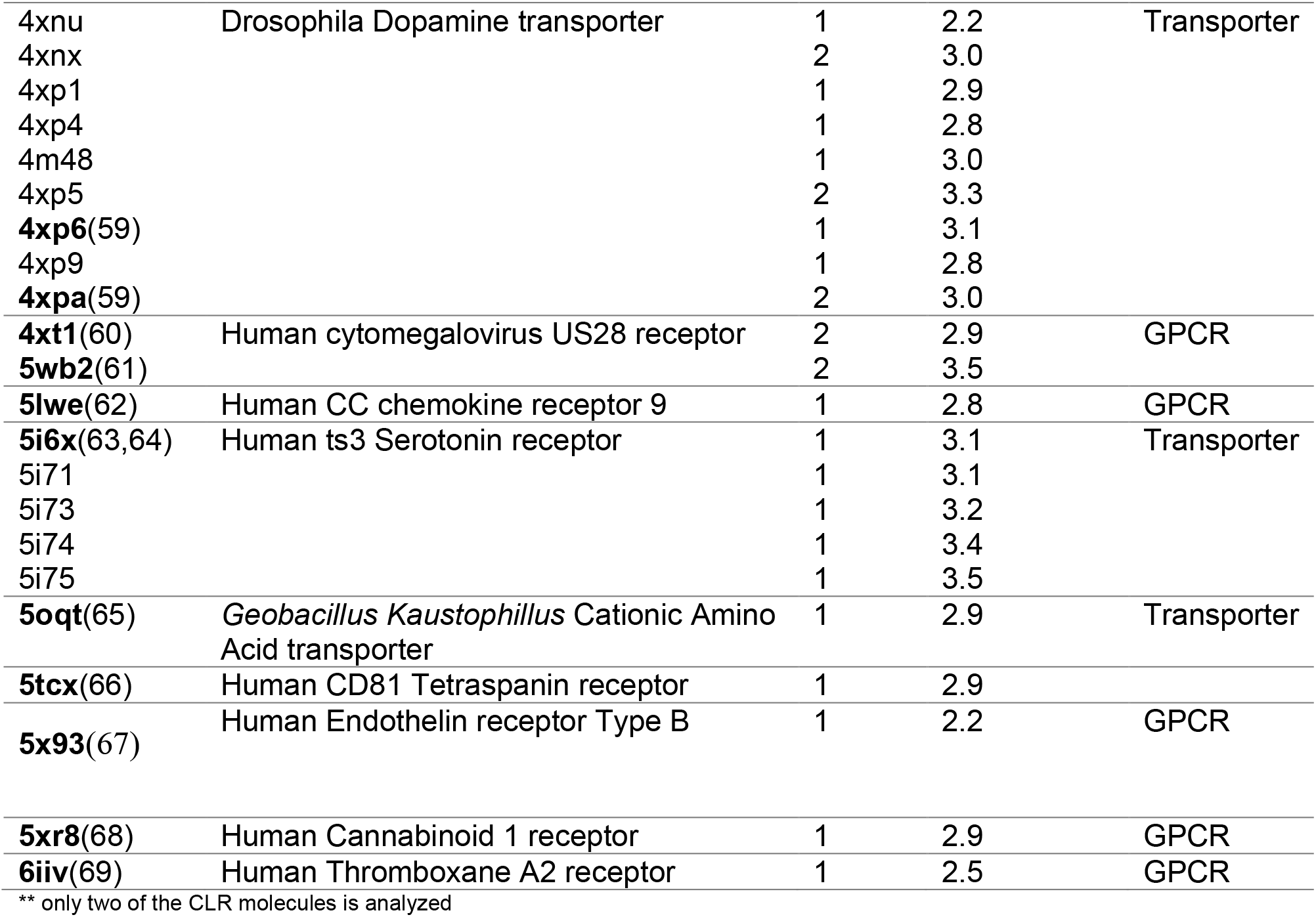
Summary of Cholesterol-IMP Dataset

### The CRAC and CARC motifs failed to align with CLR-IMP interactions in this dataset

The CRAC motif occurred 99 times and the CARC motif occurred 174 times in our CLR-IMP dataset. 65% of the motifs were in the lipid exposed transmembrane regions of the IMPs. From the co-crystallized CLRs 69% (29) of them were within 6 Å of either a total or partial CRAC and CARC motifs. Fantini et al. established that CLR recognition requires the CRAC and CARC motif to have a minimum of five residues(9) and for one of the residues to be a tyrosine or phenylalanine to anchor the tetracyclic rings(70) of CLR. Only nine of the 29 motifs met the minimum residue requirement and only two of the 29 motifs included an anchoring aromatic residue. The nine CRAC/CARC motifs that met the minimum residue number were all located on one transmembrane helix (TMH). In contrast, interaction sites in our dataset included two or more TMHs 82% of the time (see below). The tertiary structure of the two CRAC/CARC motifs that fulfilled the minimum residue number requirement and contained a phenylalanine of tyrosine residue are shown in Figure 2 and in Table 2. In both cases the CRAC/CARC motif residues only made contacts with the head group and rings A and B of cholesterol indicated in yellow. The interaction site depicted in the co-crystal structures (gray) contacted the entire CLR molecule.

**Table 2.** Summary of CRAC and CARC Motifs in Contact with Cholesterol

| CLR | CRAC | CARC | CRAC/CARC |
| --- | --- | --- | --- |
| Within 6 Å cutoff | 11 | 18 | 29 |
| Min. # residues | 4 | 5 | 9 |
| PHE or TYR<br>contact | 1 | 1 | 2 |

**Figure 2.**
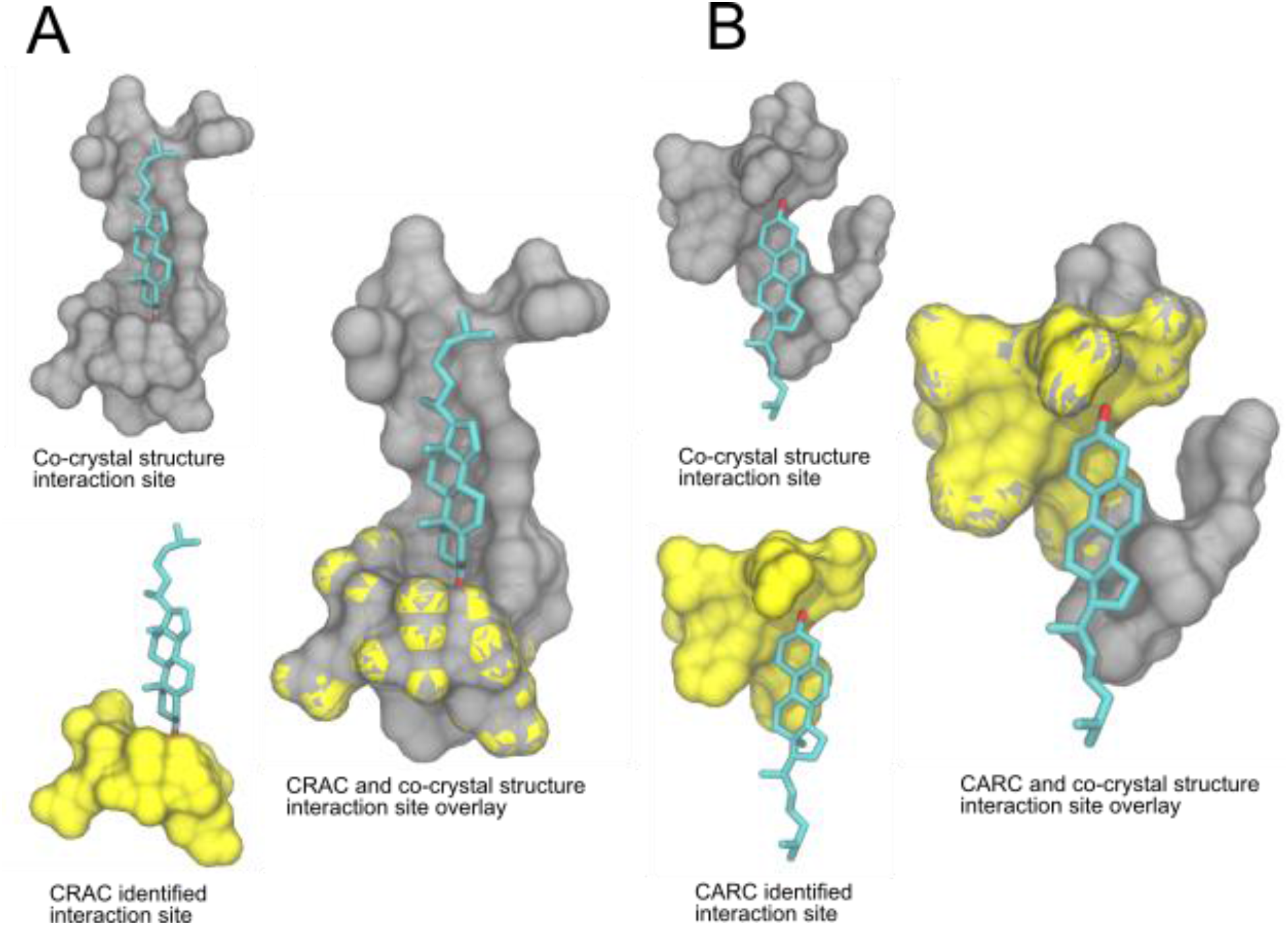
Two CRAC/CARC motifs that met the requirements defined by Fantini et al(9). A. Surface representation of the co-crystal structure of the serotonin 2A receptor bound to CLR (PDB: 6a93) in gray overlaid with the CRAC identified interaction site in yellow. B. Surface representation of the co-crystal structure of the thromboxane A_2_ receptor bound to CLR (PDB:6iiv) in gray overlaid with the CARC identified interaction site in yellow. The cholesterol molecule is in cyan.

### CLR-IMP binding sites can be clustered based on spatial arrangement of residue types

We used a combination of automated and manual techniques to cluster CLR-IMP interaction sites. The automation included derivation of interaction fingerprints that described the sites physically and chemically. The manual techniques included superposition of residue types that interacted with the tetracyclic rings, the iso-octyl tail, and the 3β-OH group of CLR. The dataset was arranged into nine distinct clusters as shown in Figure 3. Each cluster contains at least two examples of CLR-IMP binding pockets except for structure D, which is a singlet.

**Figure 3.**
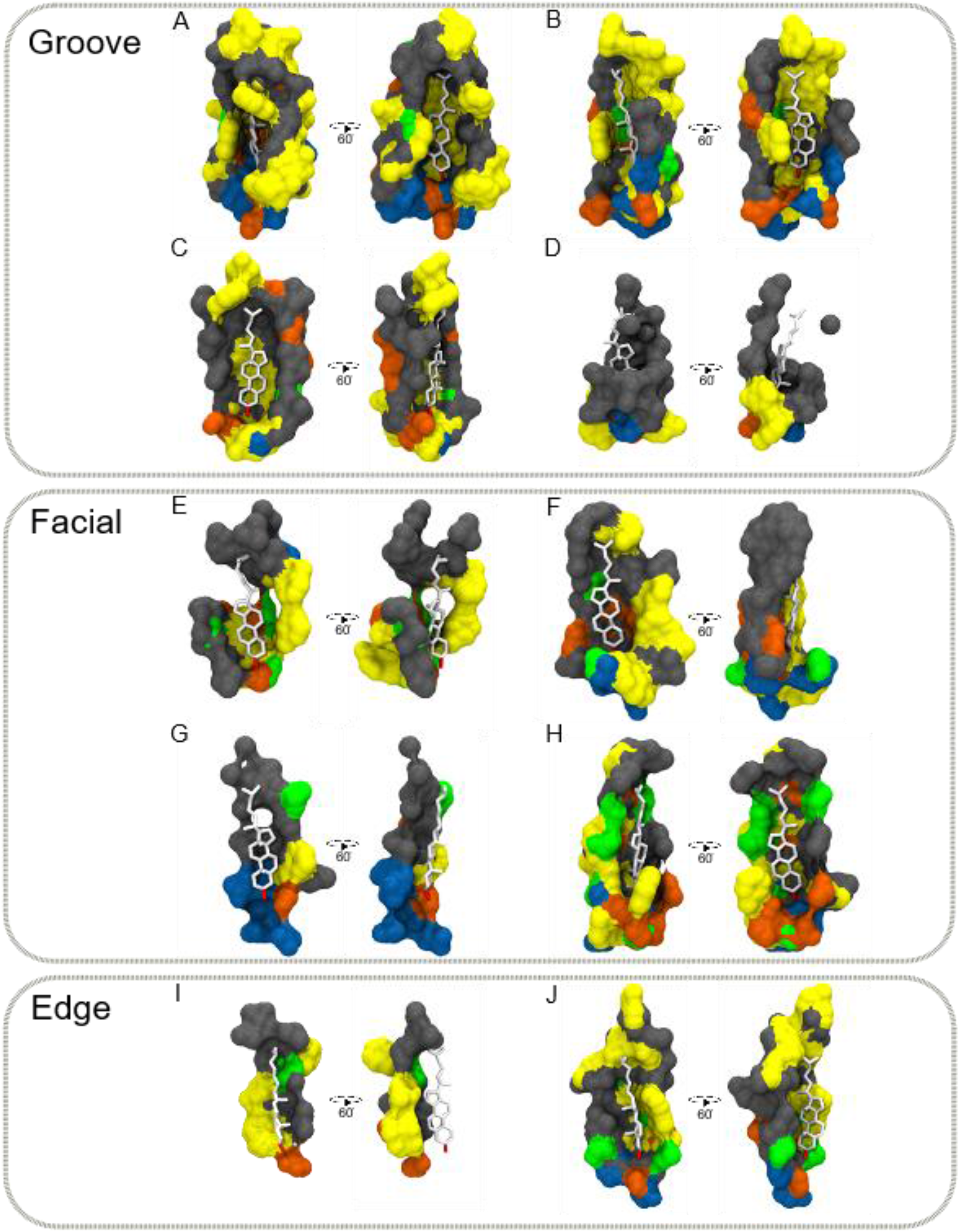
Clusters of CLR interaction sites in IMPs. For each cluster, a 3D surface representation of the sidechains surrounding the cholesterol molecule is shown and colored according to residue type. Interaction sites are categorized by pocket type: groove, facial, edge. Two views of each cluster are shown with a rotation of 60°. The number of interaction sites in each cluster is : A(9), B(4), C(7), D(1), E(3), F(5), G(2), H(4), I(2), J(7). Color key: aromatic, yellow; polar-charged, blue; polar-uncharged, orange; non-polar, gray; special (C, G, P), green; cholesterol, white.

These clusters can be grouped into several interaction site types we term groove, facial, or edge. Interaction site type groupings were based on the percent of residue types that contacted the head group, tetracyclic rings, and tail of the CLR molecule, and the number of TMHs involved. In 78% of instances grooves encompassed residues from three or four helices. The iso-octyl tail was often contacted by aromatic residues and the 3β-OH group interacted with polar residues (Figure 3). The facial sites encompassed residues from two to three helices in 88% of the instances. The iso-octyl tail interacted with non-polar residues and the 3β-OH interacted with polar and special residues (CYS, GLY, PRO). Lastly, the edge sites encompassed residues from one to two helices in 78% of instances. The iso-octyl tail interacted with non-polar residues and the 3β-OH interacted with aromatic and polar-uncharged residues. The edge site residues surround the rim of the CLR molecule. The asymmetric preference is more pronounced because the aromatic residues wrap around CLR to facilitate CH-π interactions with the α-face and the non-polar Cβ-branched residues intercalate with the β-face of CLR. In the next sections define these three sites are defined in further detail.

### Residue type preference based on the α- or β-face of CLR

For each cluster, we calculated the likelihood of observing certain residue types in contact with the six segments (3β-OH, ring A, ring B, ring C, ring D, and tail) of CLR, as defined in Figure 1. The residue type likelihoods were analyzed separately for the α- and β-faces. We grouped residues into five types based on their physical properties: ‘special’ residues are cysteine, glycine and proline; aromatic residues are tyrosine, tryptophan, and phenylalanine; non-polar residues are leucine, isoleucine, valine, methionine, and alanine; polar-uncharged residues are serine, threonine, glutamine, and asparagine; and polar-charged residues are histidine, aspartate, glutamate, arginine and lysine. We appreciate that some of the residues classified as polar and charged might be neutral in the hydrophobic environment of the membrane.

For the tetracyclic ring, we distinguish pronounced preferences (i.e. when residues of one class shows a preference for contacting one face over the other) from minute preferences of contacting one face of a particular ring (A, B, C or D) over the other. The groove sites (Figure 4) contacted the iso-octyl tail with aromatic residues and the 3β-OH group with polar-uncharged and -charged residues. Polar-uncharged residues were unlikely to contact the α face. In the case of special residues, contact likelihood was similar for both faces, but Ring C from the α face perspective was unlikely to touch residues from the special group. The aromatic residues had equal representation regardless of face.

**Figure 4.**
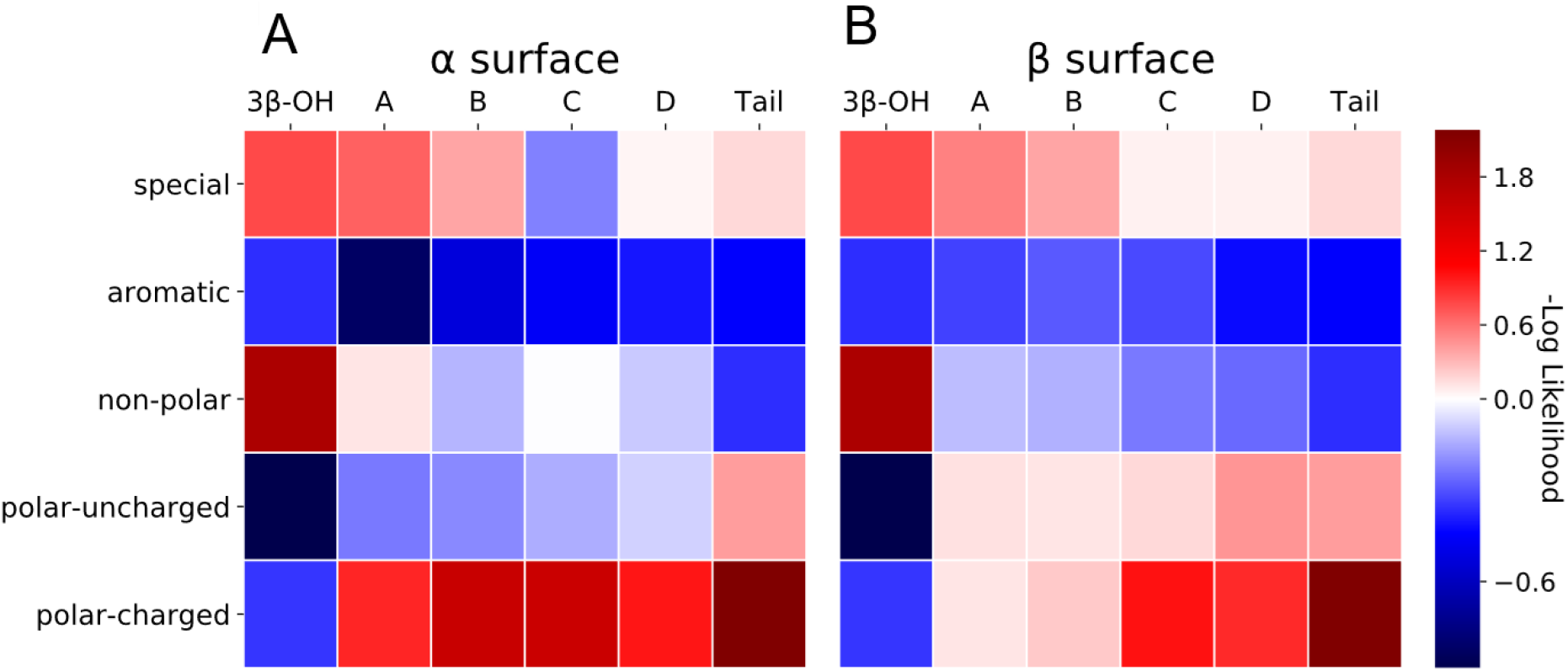
Groove interaction site residue type occurrence heatmap including 21 proteins. The color change from red to blue indicates a significant occurrence more than random. The more negative the log likelihood the more significant finding a particular residue type contacting a segment of cholesterol. Special residues are cysteine, glycine, and proline.

The facial sites (Figure 5) contacted the iso-octyl tail with non-polar residues and the 3β-OH group with special and polar uncharged residues. Special residues were unlikely to contact the β face. The more minute differences were shown in the non-polar residues where the likelihood of observed contacts was average (Ring A, B, C) or more likely (Ring D) from the β face perspective. The likelihood of aromatic residue making contacts started to decrease in the facial residues as shown in the lighter blue from the α face perspective and the complete lack of contacts at Ring A. This could be due to a decrease in TMHs involved in this type of pocket.

**Figure 5.**
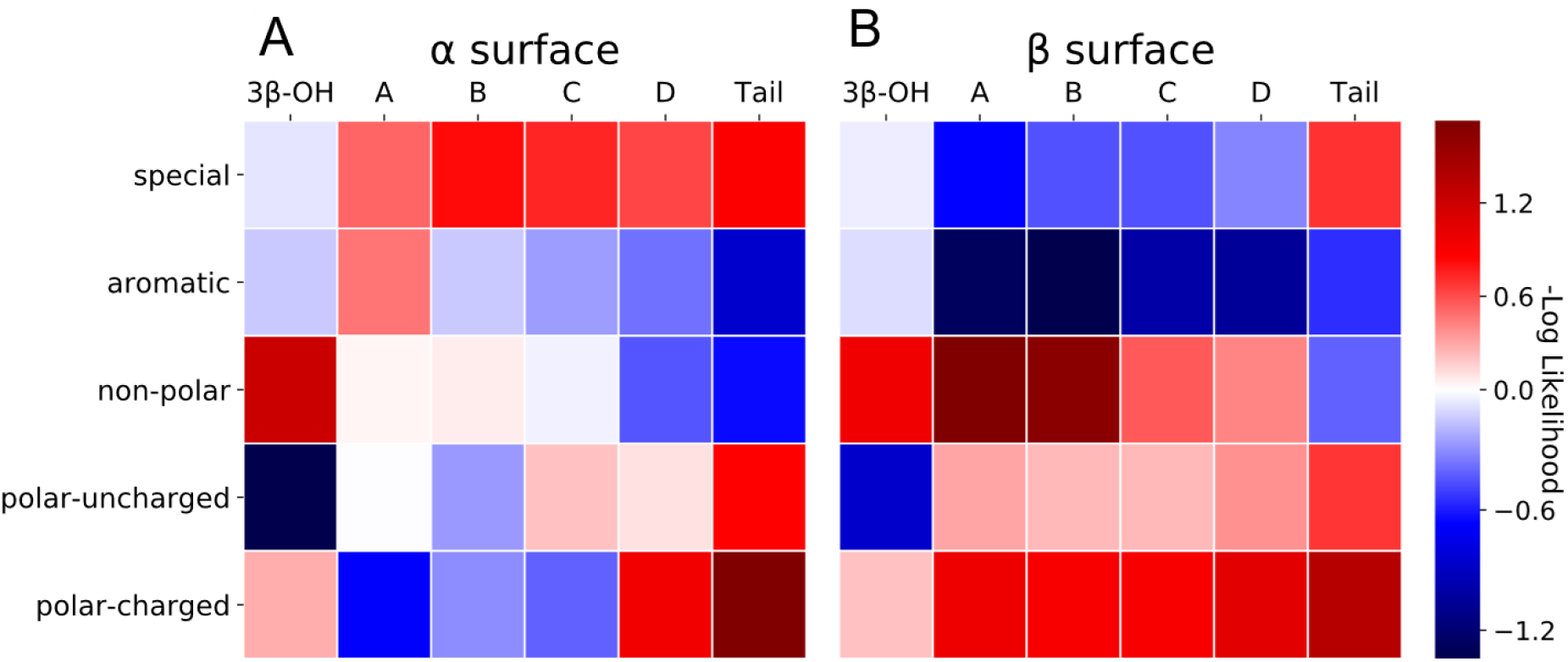
Facial interaction site residue type occurrence heatmap including 14 proteins. The color change from red to blue indicates a significant occurrence more than random. The more negative the log likelihood the more significant finding a particular residue type contacting a segment of cholesterol. Special residues are cysteine, glycine, and proline.

In edge sites (Figure 6), the iso-octyl tail contacted non-polar residues while the 3β-OH group contacted polar-uncharged and aromatic residues. In the edge sites we observed a much more pronounced degree of residue type CLR asymmetric preference. Only this type of interaction site exhibited a pattern wherein aromatic residues were biased to interact with the α face over the β face. Also, the non-polar residues exhibited the opposite preference by preferentially making contacts with the β face over the α face. In the more specific case, special residues had a bias toward making contacts with the α face of Rings A and B of CLR.

**Figure 6.**
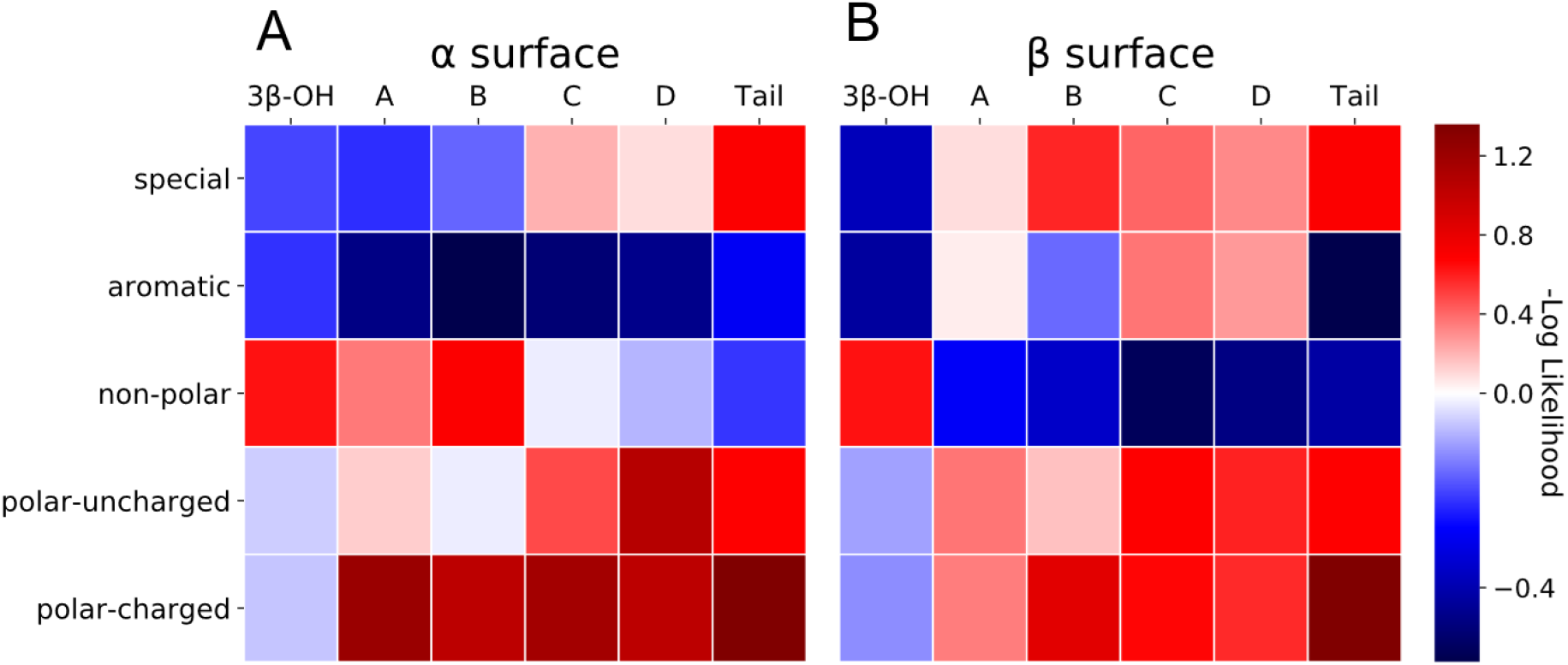
Edge interaction site residue type occurrence heatmap including 9 proteins. The color change from red to blue indicates a significant occurrence more than random. The more negative the log likelihood the more significant finding a particular residue type contacting a segment of cholesterol. Special residues are cysteine, glycine, and proline.

### Residues that are involved in specific CLR recognition are more conserved than for sites involved in nonspecific interactions

We proposed that in specific CLR-IMP interaction site residues evolve slower, i.e. are more conserved, than other surface residues of IMPs. Further, we extracted a subset of 30 CLR-IMP complexes where we found experimental or computational evidence that allowed us to categorize an interaction site as specific or nonspecific. Of these 30 CLR-IMP complexes, 14 were categorized as specific based on mutational, electrophysiology, or MD studies. The 16 interaction sites categorized as nonspecific are a default designation: Experiments were absent to support specific residue contacts being involved. The experiments only focused on how CLR content changed the membrane dynamics that affected conformations of the IMP. Thus, we expect that some of these sites might be misclassified and belong to the specific group. The remaining 14 CLR-IMP complexes out of the original 44 where classified as either not experimentally validated (nine) or putative artefact (five).

The interaction sites categorized as putative artefact were from two protein types. The prokaryotic homologue (GkApcT) of the solute 7 carrier cationic transporter (PDB: 5oqt) was crystallized with a CLR, but it is likely that this is actually a site for a hopanoid(65). Hopanoids are planar polycyclic hydrocarbons containing five rings compared with the four rings in sterols, and they have a variety of polar and nonpolar side chains(71). Regarding the metabotropic glutamate receptor 1 (mGluR1, PDB: 4or2) interaction sites it is unclear whether the dimeric bound CLRs are physiologically relevant. The crystal packing of mGluR1 captured a dimer mediated by TM1(72), but experiments done on mGluR2 and mGluR5 seem to point towards dimerization via TM4/TM5(73–75). mGluR5 and mGluR1 belong to the same group in the eight-protein family.

The average rate of evolution of the core, surface and interface of each CLR-IMP complexes in the two categories were computed (Figure 7). We determined surface versus core residues based on solvent accessibility of lipids in the membrane space. The red line in Figure 7 denotes the median rate of evolution of the non CLR interacting surface residues. A median rate of evolution below the red line correlates to stronger conservation. 98% of the specific CLR interaction sites and 79% of the nonspecific CLR interaction sites had median rate of evolutions significantly below the red line. A Wilcoxon Signed-Ranks sum test indicated that the median rate of evolution of the CLR interaction site residues was statistically significantly lower than the median rate of evolution of the surface residues (specific: p < 10^−28^, nonspecific: p < 0.001)(76,77).

**Figure 7.**
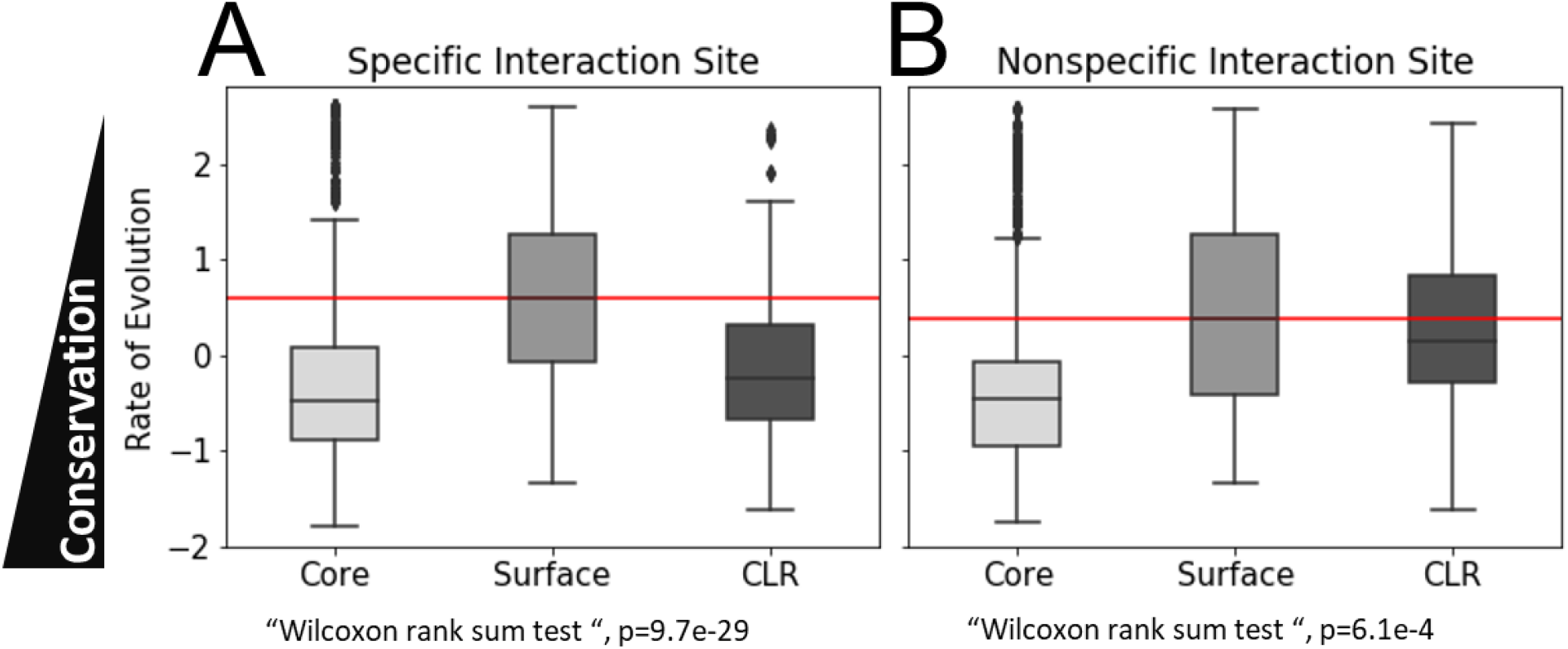
Rate of evolution calculation for residues in the core, on the surface and in the CLR interaction site. A. Specific CLR-IMP interaction sites found in Table 1 that have been experimentally validated as specific sites on the surface of those proteins. B. Nonspecific CLR-IMP interaction sites found in Table 1 that have been experimentally validated for being involved in membrane modulation. The red line denotes the median rate of evolution of the surface residues. A CLR median below the redline means it evolves more slowly than other surface residues. The Wilcoxon rank sum test p values are referring to the CLR residues versus the surface residues.

### CLR-SP interaction sites are more conserved and do not require aromatic residues

To form a baseline for specific and nonspecific CLR-IMP interaction sites we investigated the differences in tertiary structure of CLR-SP and-IMP interactions. The CLR interaction site conformation of SPs form a cavity, residues completely surround the CLR molecule, while IMPs tend to make contacts with one face of CLR. According to previous work there is an increase in polar and/or charged residues in the CLR interaction sites of SPs(78). Oppositely, in our analysis the presence of aromatic residues was almost non-existent (5%) in the SP interaction sites. While they made up 35% of IMP interaction sites. The drastic decrease in aromatic residues in SP interaction sites could be due to the absence of a membrane environment where the aromatic residues are used to anchor CLR from the non-annular lipid space.

Additionally, the average rate of evolution of the core, surface and interface was calculated for the ten CLR-SPs (see Supplementary Figure 1). Most of these proteins act in the CLR homeostatic pathway so we expected the rate of evolution of the interface residues to be similar to the core and have a large effect size. Effect size is the difference between the average rate of evolution of the residues involved in the CLR interaction site and the average rate of evolution of the other surface residues. Every protein except the ORP1-ORD complex (PDB: 5zm5) had some experimental and/or MD simulation studies that confirmed CLR as a functional or allosteric modulator. All residues involved in CLR binding in the SPs had rates of evolution lower than surface residues. These conclusions are based on a ten protein dataset (Supplementary Table 1), so sample size must be considered, but outcomes are consistent with Ounjian et al(78).

### Classification of interaction site type based on experimentally derived effect sizes

Due to the correlation between a residues rate of evolution and its presence in an interaction site, we matched the interaction site tertiary arrangement (groove, facial, edge) to conservation. for this purpose, we created a conservation scale to rank specific and nonspecific CLR interaction sites as shown in Figure 8A. The scale was based on the difference between the mean rate of evolution of the surface residues and the residues involved in the CLR interaction site, also known as the effect size. The more negative the effect size the greater the difference in conservation of the CLR interaction site residues and the surface residues. The specific interaction site effect size ranges between −1.1 and −0.7. The effect size of the CLR-SP dataset(79) (Supplementary Table 1) was also calculated to compare the specific interaction sites to known specific CLR-SP interactions as a baseline comparison. Conserved CLR-SP interaction site effect sizes were between −1.4 and −0.8. Nonspecific interaction site effect size range was from −0.6 to 0.0. There was a large variance in the effect size range of the nonspecific interaction site residues as well as an overlap with the effect size range of the specific interaction sites. This is most likely due to the experimental evidence only suggesting these IMPs are affected by the overall change of CLR in the membrane, but not directly investigating specific residue contacts. Some of these IMPs will need to be reclassified as making specific CLR interactions when more experimental data becomes available. By contrast, The CLR-rhodopsin interaction (PDB: 3am6) was labelled as nonspecific because multiple studies supported the notion that modification of the membrane environment is the driving factor for its preference for the inactive state(80–82). The mean effect size for both interaction sites was −0.13.

**Figure 8.**
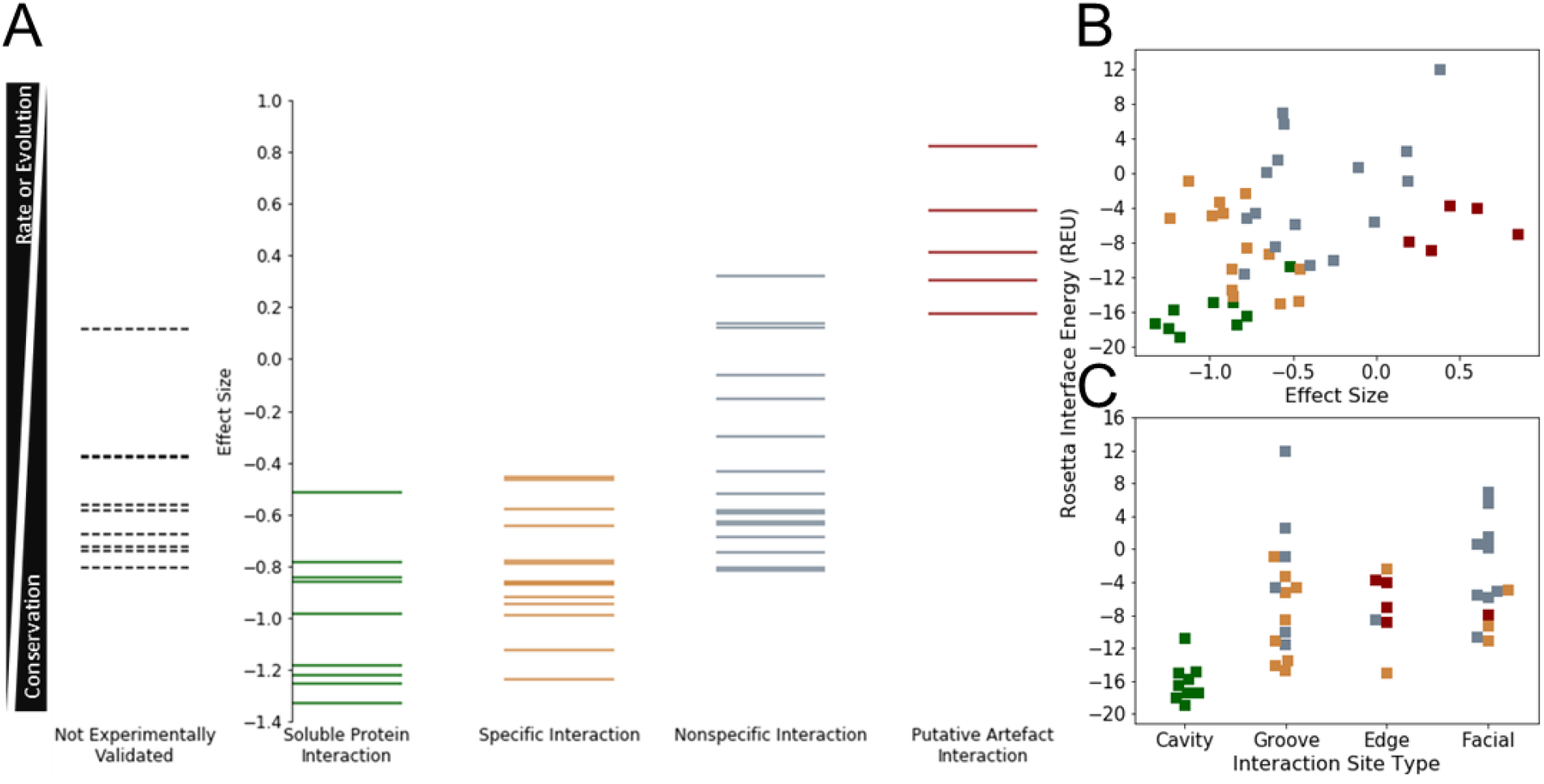
A. Conservation Scale. Each column represents the CLR interaction site categories shown in Supplementary Figure 1. Right of the axes is each of the 35 interaction sites ranked based on the biochemical/MD simulation data Left of the axes are the nine interaction sites that had no biochemical/MD to support their categorization. Sites were ranked based on their effect size. B. Rosetta interface energy versus effect size plot for CLR interaction site categories. C. Rosetta interface energy versus interaction site type plot for CLR interaction site categories. The color of the squares correspond to interaction site types ranked in A. Effect Size = (X̄(CLR rate of evolution) − X̄(Surface rate of evolution)

### Rosetta Interaction Energy differences for specific and nonspecific CLR interactions

The Rosetta Macromolecular Software Suite(83,84) was used to score CLR interaction sites amongst four groups (specific, nonspecific, soluble, and putative artefact). In Figure 8B we compare the Rosetta Interaction Energy (RIE) to the effect size and interaction site. The four groups were able to accumulate into distinct groups with some overlap between the specific- and nonspecific-IMP sites. The CLR-SP interaction site effect size correlated with the most negative RIE (−16.07 REU) as expected. The specific sites group average RIE was −8.51 REU and always less than zero. The nonspecific group was more variable with an average RIE of 0.135 REU. The average RIE of the putative artefact sites was −6.34 REU with their effect sizes greater than zero. Additionally, 80% of the specific interaction sites in the 30 CLR-IMP subset were identified as groove sites, while 62% of the nonspecific interaction sites were facial. The edge sites were equally represented in both (shown in Figure 8Figure 8 A. Conservation Scale. Each column represents the CLR interaction site categories shown in Supplementary Figure 1. Right of the axes is each of the 35 interaction sites ranked based on the biochemical/MD simulation data Left of the axes are the nine interaction sites that had no biochemical/MD to support their categorization. Sites were ranked based on their effect size. B. Rosetta interface energy versus effect size plot for CLR interaction site categories. C. Rosetta interface energy versus interaction site type plot for CLR interaction site categories. The color of the squares correspond to interaction site types ranked in A. Effect Size = (X̄(CLR rate of evolution) − X̄(Surface rate of evolution) C).

## DISCUSSION

IMP structure and function can be affected by the presence of CLR as we expressed in the introduction. However, the underlying molecular mechanisms remain difficult to parse within a varying set of protein types. The major reason for this is that it is near impossible to separate specific and nonspecific effects in a cooperative assembly like membranes. The categorization of CLR-IMP interaction sites allows for increased identification of sterols in low-resolution density cryo-electron microscopy structures and allosteric sites that effect orthosteric ligands in cases of the type-1 cannabinoid receptor(85) (PDB: 5xr8) and the translocator protein in PET imaging(86,87). To computationally identify and rank CLR interaction sites we need to determine the necessary statistics to define specific and nonspecific sites.

The sequence-based CRAC/CARC motifs are accepted in most cases as enough to describe specific CLR-IMP interaction sites. In our dataset the motifs were neither sufficient nor necessary to identify CLR-IMP interaction sites. We identified 273 CRAC and CARC motifs, but 99% of them were insignificant in capturing the CLR-IMP interaction sites in our dataset. For those motifs that did contribute residues to CLR-IMP interaction sites, there were only two examples (PDB: 6iiv and 6a93) where all the key residues were engaged in a CLR interaction. We classified the CLR-serotonin 2A receptor (PDB: 6iiv) as nonspecific and the CLR-thromboxane A_2_ receptor (PDB: 6a93) as a specific interaction site as derived from sequence conservation and tertiary structure of the CLR-IMP interaction. We were unable to identify a repeatable CRAC/CARC motif pattern that classified specific versus nonspecific CLR-IMP interactions sites. Instead, 89% of the CARC/CRAC motifs that were within 6 Å of CLR, donated a residue here or there to interaction sites, but not in a consistent fashion (see Figure 2 and Table 2). Due to a lack of consistency in CRAC/CARC motifs we delved into characterizing our dataset structurally and evolutionarily. The interaction site type clusters (groove, facial, and edge) allowed us to delineate similar binding modes across IMP type and determine conformations that maybe necessary for specific CLR-IMP interaction sites. Specific sites as determined by rate of evolution were characterized by grooves with a RIE average of −5.45 REU, while nonspecific sites were characterized by facial interactions with a RIE average of - 4.67 REU.

Stereo-specificity of CLR-IMP interactions has become even more important in the investigations(34) This study further points out how these minute differences play a large role in recognition. 73% of the 44 CLR-IMP interactions sites showed an explicit asymmetrical preference. There is 14 α facing interaction sites and 16 β facing ones. In 28 of the IMP structures used in the study 13 we co-crystallized with two or more CLR molecules. The CLR molecules were within 6 Å of each other in seven of them. In this small subset we noticed that in five of the CLR molecule pairs the IMP interacted with either the α- or β-face of one CLR and both faces in the other. We speculate that in some instances that multiple CLR recruitment at an interaction site is signaled by the asymmetric binding event of one CLR molecule. Studies have shown that direct and indirect CLR interactions with GPCRs can cooperatively (positively or negatively) effect oligomerization(88), stability(89), and the binding of orthosteric ligands(90– 92). Multiple CLR molecules are usually found bound to the IMP. Due to the snapshot nature of crystal structures we are unable to determine which CLR bound first, but with MD simulations we can further investigate.

In the future, distinction of specific from nonspecific CLR-IMP interactions can be improved e.g. incorporating MD data such as residence time to complement some of the inconclusive experimental validation results. In the future we plan to further differentiate specific and nonspecific by using MD simulations.

Computational design has the potential to provide a general, complementary approach for CLR recognition in which design features and specificity can be rationally programmed. The increase in co-crystal CLR-IMP structures will enable development of statistical potentials derived from inter-residue, CLR segment-residue, and CLR segment-secondary. A specificity potential can be formed based on the categorization of the interaction sites based on their tertiary structure as one of the three types of clusters (groove, facial, and edge) and the calculation of the rate of evolution effect size. Once the energy of all atomic interactions is calculated with the current Rosetta score function the specificity potential will be used to determine if a site is more likely to be specific or nonspecific.

In conclusion, the structural and evolutionary statistics gained from this study will allow to set the foundation for better ranking/differentiation of possible specific or nonspecific CLR-IMP interaction sites. This information will be implemented into the RosettaLigand(93) scoring function for docking and design of specific CLR-IMP interactions.

## MATERIALS AND METHODS

### Dataset compilation

At the time of writing the PDB^6^ contains 120 high-resolution crystal structures of CLR-IMP complexes. A dataset of 44 CLR-IMP (Table 1) interaction sites were investigated to understand CLR recognition. All CLR-IMP complexes were determined through X-ray crystallography with a resolution of 3.5 Å or better. Structures were chosen based on either preliminary experimental evidence that suggested a functional effect or if no evidence was given that CLR was added to aid crystallization.

### Rosetta Interaction Energy Score Protocol

We adapted the MPRelax application(94) to use on the 44 CLR-IMP co-crystal complexes to refine the structure and optimize the membrane positioning and orientation. The CLR-IMP interface score was calculated with a modified version of the Rosetta InterfaceScoreCalculator Mover with the option added to score the CLR ligand within the membrane. After calculating the Rosetta score for the bound CLR-IMP complex, IMP and CLR are moved away from each other along the XY plane of the membrane and scored separately. The interface score is calculated as difference between the score values IMP and CLR in the bound and unbound state. The protocol capture is provided in the Supporting Materials.

### CLR-IMP residue interactions

We first used a 2D visualization of general hydrogen bonds and hydrophobic residue interactions was used to classify CLR-IMP interaction sites first using LigPlot^80^. Ligplot results provided a starting point to map residue contacts but failed to account for the asymmetric characterization of CLR binding pockets. Then we wrote a script to calculate residue contacts and potential hydrogen bond formation of each interaction site within 6 Å. CLR was subdivided into six segments (see Figure 1) to categorize how often certain residues interact with these motifs and if the α- or β-face of the CLR molecule is preferred‥ Results were divided by residue type: aromatic, polar-charged, polar-uncharged, non-polar, or special (CYS, GLY, PRO). Percent interaction of each residue type with certain CLR motifs were normalized and converted into a likelihood metric.

### Clustering techniques to determine CLR binding pocket descriptions

CLR-IMP interaction sites were manually superimposed based on the placement of the CLR molecule in Pymol(95). Residues and backbone atoms of the IMP within 6 Å of the respective CLRs were aligned. Using the quantified IMP interactions from python we selected ten unique clusters containing all 44 CLR-IMP complexes. Manual clustering was based primarily on aromatic residues contacting one of the tetracyclic rings and then residue types surrounding the tail or head group.

CLR-IMP interaction fingerprints were constructed based on Fergus Boyles *et al’s* workflow(96). We combined ligand- and structure-based features. The ligand features were computed only once since CLR was the only ligand. The ligand-based features used 200 molecular descriptors for each ligand based on the Descriptors module of the Python RDKit package version 2018.03 (https://www.rdkit.org/, accessed May 17, 2019). The descriptors are conformation independent and are categorized as experimental properties or theoretical descriptors. Any features with zero variance or null valued across the dataset were excluded. The structure-based features were computed using the machine learning RF-Score function(97) implemented through Open Drug Discovery Toolkit (ODDT) version 0.6(98).

### Determine CRAC/CARC on IMP

All possible CRAC and CARC motifs were identified on each of the 28 IMPs in the dataset. The CRAC/CARC motif patterns described in the introduction were searched for based on the fasta sequence. Then we checked if any of the 44 CLRs in the dataset were within 6 Å of a CRAC or CARC motifs of the PDB. To calculate percent occurrences, we classified the areas where there was no CLR or motif found. The relative solvent accessibility(99) was calculated on all 44 interaction sites to determine this.

### Determine Core and Surface Residues using Lipid accessibility analysis

IMP residues were divided into core and surface regions based on their accessibility to lipids. We used the NetSurfP-2.0(99) server to compute relative solvent accessibility (RSA) to differentiate between the two categories. RSA was computed for each residue based on the ratio between the absolute solvent accessibility (ASA) observed in the native structure and in an extended tripeptide conformation (A-X-A). The Rosetta mp_lipid_acc mover(100) was used to specify only surface residues that interacted with the lipid environment. To identify lipid exposed residues, they use a 2D concave hull algorithm to determine water and lipid accessible boundaries. The subset of surface residues that interacted with CLR were labelled as CLR in Figure 7.

### Determine site-specific rate of evolution of CLR binding pockets

Site-specific rate of evolution was estimated using the Rate4Site(101) algorithm. The original dataset was reduced from 44 to 30 CLR-IMP interaction sites based on the availability of experimental and/or computational data that validate CLR-IMP interactions as either specific or nonspecific. A multiple sequence alignment (MSA) of each of the IMPs was used as input. MSAs were obtained by running HHblits(102) against the uniport 20 sequence database(103), with minimum coverage with master sequence set to 25%, minimum sequence identity with master sequence set to 15%, target diversity of MSA set to 5, number of iterations set to two, and E-value cutoff for inclusion set to 0.001. The output MSA was limited to a cutoff of 300 aligned sequences before being input into the Rate4Site program.

## AUTHOR CONTRIBUTIONS

B.M. and J.M. designed research. B.M. performed research, created figures and tables. B.M., B.L. and G.K. created code to analyze results. B.M., C.S., and J.M. analyzed and discussed the results. B.M. and J.M. contributed to writing of the article. All authors reviewed and edited the manuscript.

## ACKNOWLEDGEMENTS

We thank Julia Koehler Leman, Benjamin K. Mueller, Renã Robinson, Jarrod Smith, and Charles Manning for helpful discussions. B.M. was supported by National Science Foundation Graduate Research Fellowship Program awards DGE-1445197 and DGE-1937963. The research was supported through NIH NIGMS R01 GM080403 and NIH NIHL R01 HL122010. J.M. acknowledges funding by the Deutsche Forschungsgemeinschaft (DFG, German Research Foundation) through SFB1423, project number 421152132.

## SUPPORTING CITATIONS

References (104-116) appear in the Supporting Material

